# Metallothionein-2A protect Cardiomyocytes from Ischemia/Reperfusion through inhibiting p38

**DOI:** 10.1101/2021.09.09.459560

**Authors:** Ying Zhao, Xiaoli Huang, Yuanlin Lei, Jike Li

## Abstract

The resumption of coronary artery blood supply is often accompanied by myocardial ischemia/reperfusion (I/R) injury after the occurrence of myocardial infarction, shock, cardiac surgery and other events. Metallothionein-2A (MT2A) has the functions of scavenging free radicals, anti-oxidative stress, anti-apoptosis, anti-autophagy, promoting vascular growth. The activation of p38 MAPK pathway can induce cardiomyocyte apoptosis in H9c2 cardiomyocytes during I/R, thereby aggravating the myocardial I/R injury. However, it is not clear that the effect of MT2A on p38 in cardiomyocytes under I/R. A simulated I/R model was used. Our objective was to investigate the protective effect of MT2A on I/R-caused mortality in H9c2 cardiomyocytes through its influence on p38, as well as the relationships among these processes. The results indicate that both endogenously overexpressed MT2A and exogenously added MT2A can inhibit the active expression of p38 during I/R. Based on these results, I/R induces apoptosis and p-p38 in cardiomyocytes. MT2A can inhibit the active expression of p38. MT2A protects cardiomyocytes from I/R injury, and that p38 is one of the molecules of MT2A against I/R injury in cardiomyocytes.

## Introduction

Ischemic heart disease is one of the leading causes of death in human and timely reperfusion is the most effective method for the treatment of acute myocardial infarction [1]. However, studies have found that reperfusion may cause cardiomyocyte injury and this process is often called myocardial ischemia/reperfusion (I/R) injury [2]. Many factors can lead to myocardial I/R injury and oxidative stress caused by reactive oxygen species (ROS) explosion during reperfusion is a key trigger factor of myocardial I/R injury [3].

The p38 is a member of the mitogen-activated protein kinases (MAPKs) subfamily, playing an indispensable role in cell signal transduction, which can cause inflammation, cell growth, apoptosis and other biological effects [4, 5]. With the increase of ROS, p38-MAPKs signaling pathway [6] can be activated and then autophagy and apoptosis can be induced [7].

Metallothioneins (MTs) are low molecular weight proteins that bind to metal ions such as Zn and Cd [8]. MT2A, the most widely expressed member of the human MT family, exists in almost all types of soft tissue [9, 10]. MT2A is a small molecule protein consist with 62 amino acids and containing 30 percent of the residues are cysteines [11]. It is anti-inflammatory, anti-endotoxin, regulates tumor growth, maintains the homeostasis of zinc in cells, and protects cells from heavy metals [12, 13, 14]. In addition, MT2A has the functions of scavenging free radicals, anti-oxidative stress, anti-apoptosis, anti-autophagy, promoting vascular growth, and insulin resistance in adipocytes [11, 15, 16].

It was found that p38 signaling pathway is closely related to the occurrence and development of myocardial I/R injury [17]. MT2A may protect cardiomyocytes from I/R injury through p38 signaling pathway. To confirm this hypothesis, H9c2 cardiomyocytes were subjected to I/R in vitro. The purpose of this study was to explore the protective effect of MT2A on cardiomyocytes during the process of I/R through p38 signal pathway.

## Materials and methods

### Reagents and materials

Dulbecco’s modified Eagle’s growth medium (DMEM), fetal bovine serum (FBS) and trypsin were purchased from Biological Industries (BioReagent, Israel). DMSO was obtained from Sigma Aldrich (St.Louis, MO, United States). The p38 primary antibody, horseradish peroxidase-conjugated secondary antibodies and Zn7-MT2A protein were acquired from Santa Cruz Biotechnology (Santa Cruz, CA, United States). Caspase-3 and α-Tubulin primary antibodies were bought from Abcam (Cambridge, United Kingdom). BCA protein assay kit and 2-(4-Amidinophenyl)-6-indolecarbamidine dihydrochloride (DAPI) Staining Solution were obtained from Beyotime Biotechnology (Beijing, China). All chemical reagents were of at least analytical grade.

### Cell line and culture

The H9c2 (2-1) cell line is derived from rat embryonic cardiomyocytes from the ATCC cell bank. Cardiomyocytes were cultured in DMEM containing 1% penicillin, streptomycin and 10% fetal bovine serum, and then the cells were incubated in a humidified incubator at 37°C, 21% O_2_ and 5% CO_2_. The medium was replaced every 2-3 days, and cells were subcultured or subjected to experimental procedures at 80-90 % confluence.

### Ischemia/Reperfusion

According to previous literature reports, hypoxia control chambers were used to construct ischemia model and I/R model ^[18]^. The control cells were cultured in an environment containing 21% O_2_, 5% CO_2_ and 74% N_2_. Ischemia cells were cultured for 5 hours in an anoxic environment containing 1% O_2_, 5% CO_2_, and 94% N_2_. I/R cells were reperfused for 1 hour after ischemia for 5 hours under conditions of 21% O_2_ and 5% CO_2_.

### Morphological study under fluorescence microscope

#### Morphological study of unstained cells

The unstained cells were observed by fluorescence microscopy (white light) after cultivation.

### DAPI staining

The H9c2 cardiomyocytes in the logarithmic growth phase will be made into cell suspensions. The cell density was adjusted to 5 × 10^5^ cells/ml and seeded in a 6-well culture plate at 1 ml/well. The cells were cultured in an incubator at 37°C and 5% CO_2_ for 48 hours. After the cells are completely attached, ischemia and I/R were treated respectively. The control cells were only added to the same volume of culture. After the cells were processed, they were rinsed 3 times with PBS. The cells were stained for 10 minutes at room temperature with DAPI working solution. Finally, they were viewed and photographed under a fluorescence microscope (Leica Biosystems, Germany).

### Lentiviral production and transfection

MT2A overexpresses lentiviral vector pLent-EF1a-MT2A-FH-CMV-GFP-P2A-Puro (LV-MT2A) and control vector pLent-EF1a-FH-CMV-GFP-P2A-Puro (LV-GFP) were constructed in cooperation with ViGene Biosciences.inc (Shandong, China). The sequences of the MT2A primers used were as follows: Forward, 5′-CAGATGGATCC-TGCTCCTGC-3′ and reverse, 5′-CTCTTTGCAGATGCAGCCCT-3′. The cloned gene was confirmed by sequencing (CMV promoter primer, 5′ -CGCAAATGGGCGGTAG-GCGTG -3′). The titer of LV-MT2A virus is 1.0 × 10^8^ TU/ml. The titer of LV-GFP virus is 3.0 × 10^8^ TU/ml.

The cells were seeded in a six-well plate at 1 × 10^6^ cells/ml. The culture solution was aspirated the next day and then washed twice with PBS. Transfection was performed at 10^4^, 10^3^, 10^2^, 10 for multiplicity of infection (MOI). DMEM containing 10% FBS was added after 4h. Change the fresh medium after 24h. Since LV-MT2A virus and LV-GFP virus carry green fluorescent protein, the number of cardiomyocytes producing green fluorescence can be observed by fluorescence microscopy (Leica Biosystems, Germany) at 24h, 48h and 72h after virus transfection of H9c2 cardiomyocytes. We randomly selected 5 non-repeating areas and calculated the transfection efficiency. The titer lentivirus and transfection time curves were plotted as mean values. Transfection efficiency = (expression of green fluorescent cardiomyocytes/total number of cardiomyocytes) × 100%.

### Western blot analysis

Add 2 ml of phosphate at 4°C to the flask and scrape off the cells. Cells were lysed with Phenylmethanesulfonyl fluoride buffer for 30 min and centrifuged at 12, 000 × g for 15 min prior to supernatant collection. The protein concentration was quantified using the BCA protein assay kit. The sample was separated by 12% SDS-PAGE and transferred to a polyvinylidene fluoride (PVDF) membrane. After the film was transferred, the PVDF membrane was immersed in TBS and rinsed 3 times for 10 minutes each time. The membrane was then placed in a 2% BSA blocking solution and blocked for 2 h at room temperature. The primary antibodies such as caspase-3, p-p38, and α-Tublin were diluted at a ratio of 1: 1000. The membrane was incubated with primary antibody overnight at 4°C. The PVDF membrane was washed with TBST on a horizontal shaker the next day and then incubated with horseradish peroxidase-conjugated goat anti-rabbit secondary antibody (1: 1000) for 1 hour. In a dark environment, the film was placed in a Tanon 5200 Multi fully automated chemiluminescence/fluorescence image analysis system (Shanghai Tianneng Technology Co., Ltd., Shanghai, China) and photographed. Finally, the membrane was quantified using the TanonImage image analysis program (Shanghai Tianjie Instrument Co., Ltd., Shanghai, China).

### Statistical analysis

All the experimental data were statistically analyzed by SPSS 19.0 software version for Windows (IBM corp., Armonk, NY, USA). All the results were expressed as means ± SD. One-way analysis was performed by one-way analysis of variance. The multivariate analysis was performed by multivariate analysis of variance. p<0.05 for the difference was statistically significant.

## Results

### Morphological changes of H9c2 cardiomyocytes

The number of vacuolated cells increased significantly under I/R. The cells become smaller and detach from the surrounding cells after I/R. DAPI staining also showed nuclear pyknosis (Fig. 1).

**Fig. 1.**
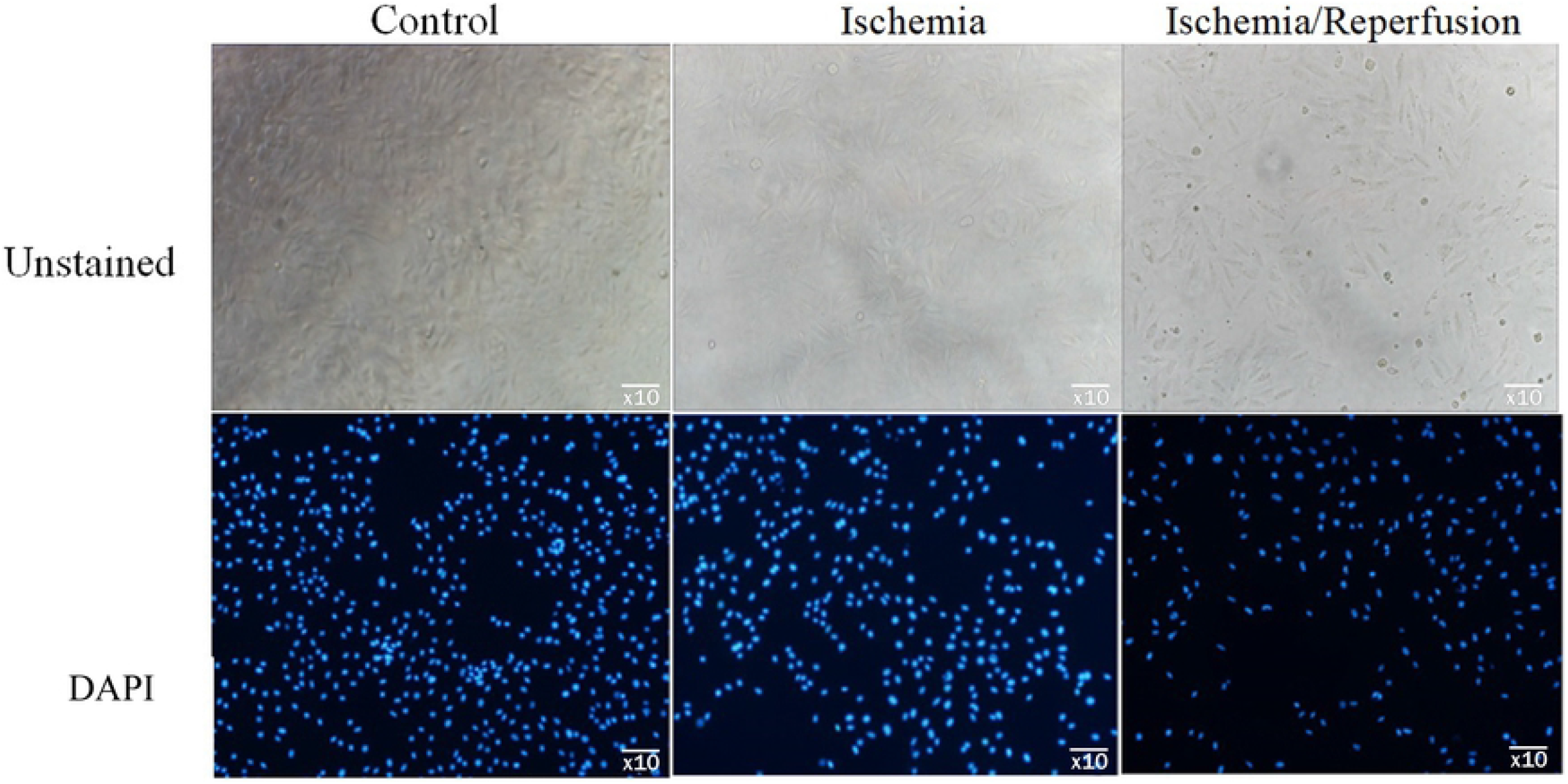
Changes of H9c2 cardiomyocytes under I/R (× 10). A-F cells were incubated under control, ischemia and I/R conditions. Images of cell morphology are shown. I/R, ischemia/reperfusion. Scale bar = 100 µM.

### Effect of I/R on expression of cleaved caspase-3

The cleaved caspase-3 and caspase-3 were detected by western blot. As shown in Figure 2, caspase-3 gradually decreased and cleaved caspase-3 increased under I/R.

**Fig. 2.**
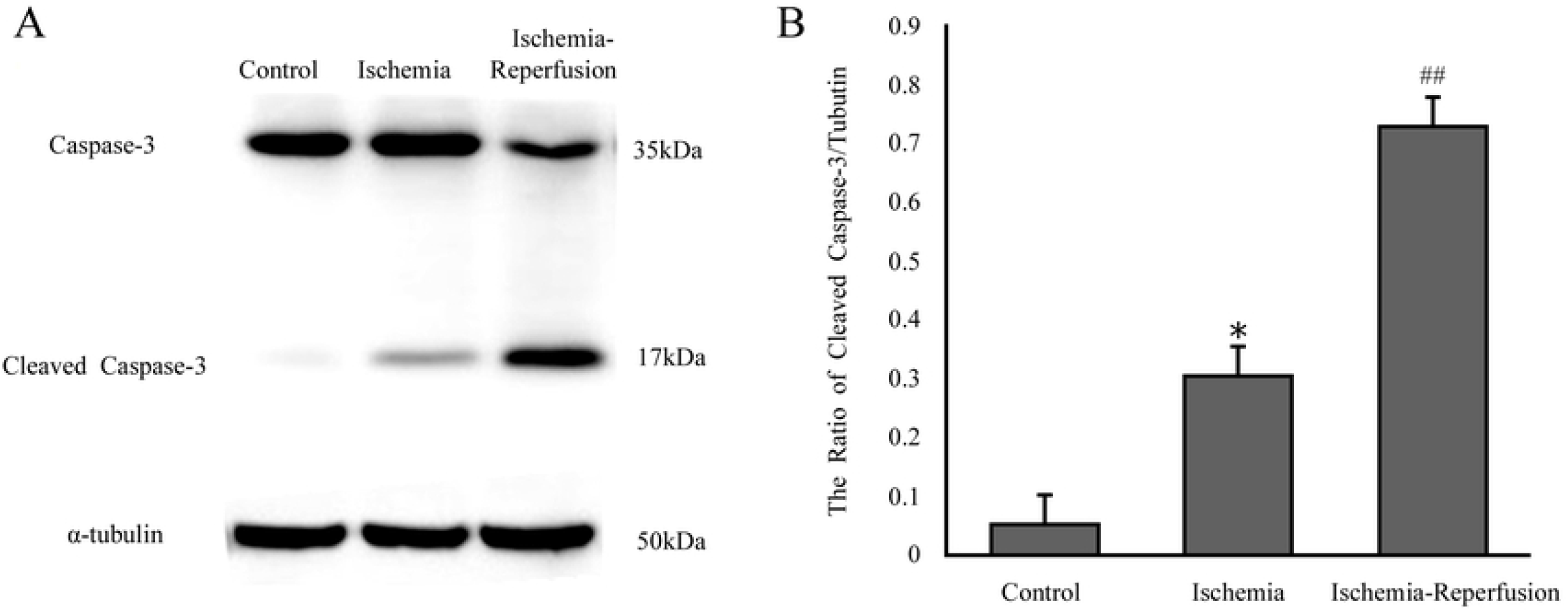
Effect of I/R on expression of caspase-3 in H9c2 cells. (A) Western blotting was used to detect cleaved caspase-3 and caspase-3. (B) quantification (n = 3). *p <0.05 vs. control group; ##p < 0.01 vs. ischemia groups. I/R, ischemia/reperfusion.

### Effects of different virus titers and transfection time on transfection efficiency of H9c2 cardiomyocytes

The transfection efficiency of LV-MT2A and LV-GFP virus decreased with the decrease of virus titer or the extension of transfection time. They reached a peak at MOI = 1000 and 48h and the transfection efficiencies were 94.4% and 95.1% respectively (Fig. 3).

**Fig. 3.**
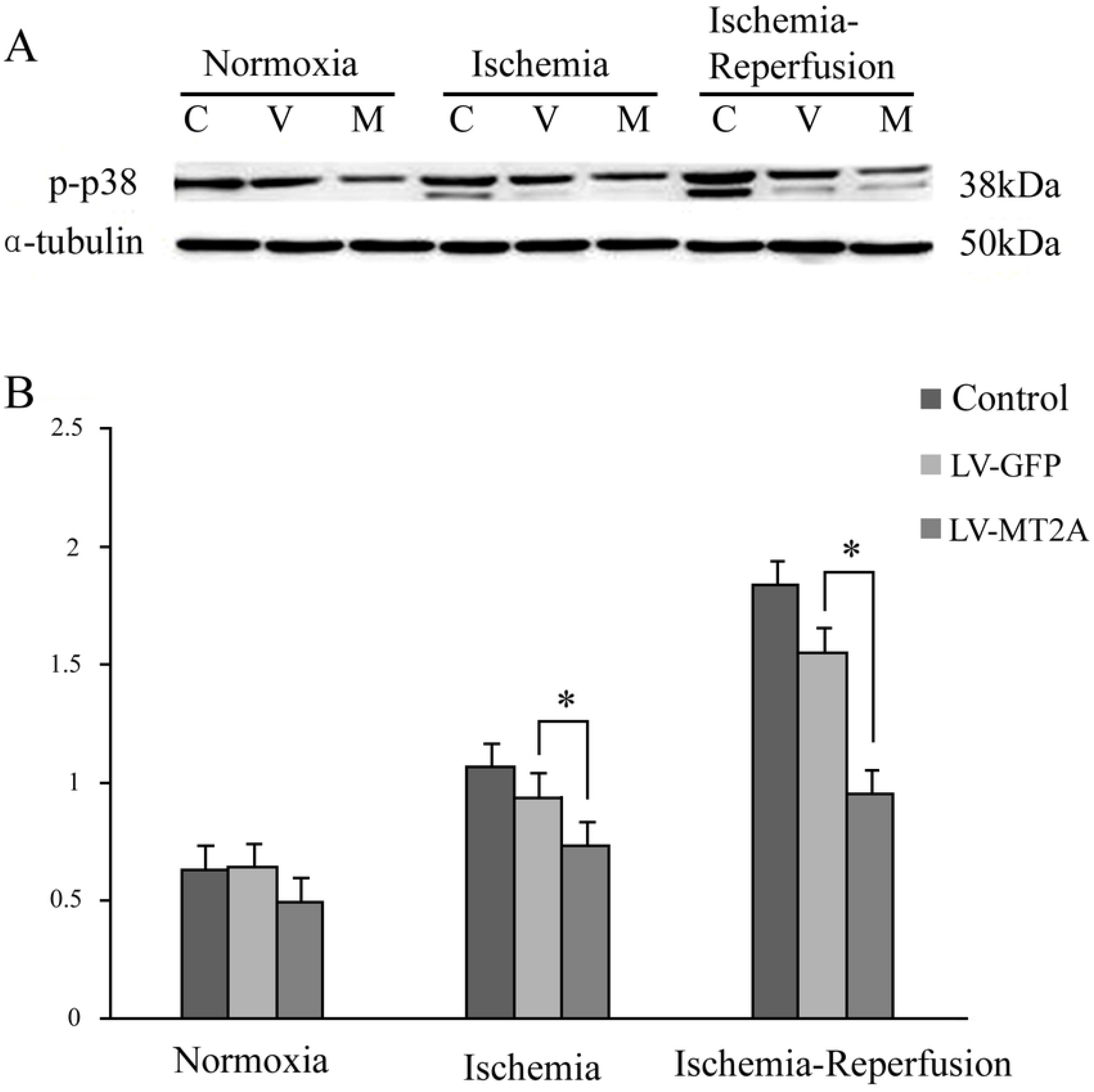
Effects of virus titer and transfection time on the transfection efficiency of H9c2 cardiomyocytes. The intensity of green fluorescence expression after virus transfection was observed under a fluorescence microscope (×10). 5 non-repetitive area radiographs were randomly selected to calculate the transfection efficiency of LV-MT2A virus and LV-GFP virus. It showed the highest transfection efficiency at MOI = 1000 and 48h in this figure.

### The expression of p-p38 can be induced under I/R condition in H9c2 cardiomyocytes

As shown in Fig. 4, the expression of p-p38 increased compared with the control group (p<0.05).

**Fig. 4.**
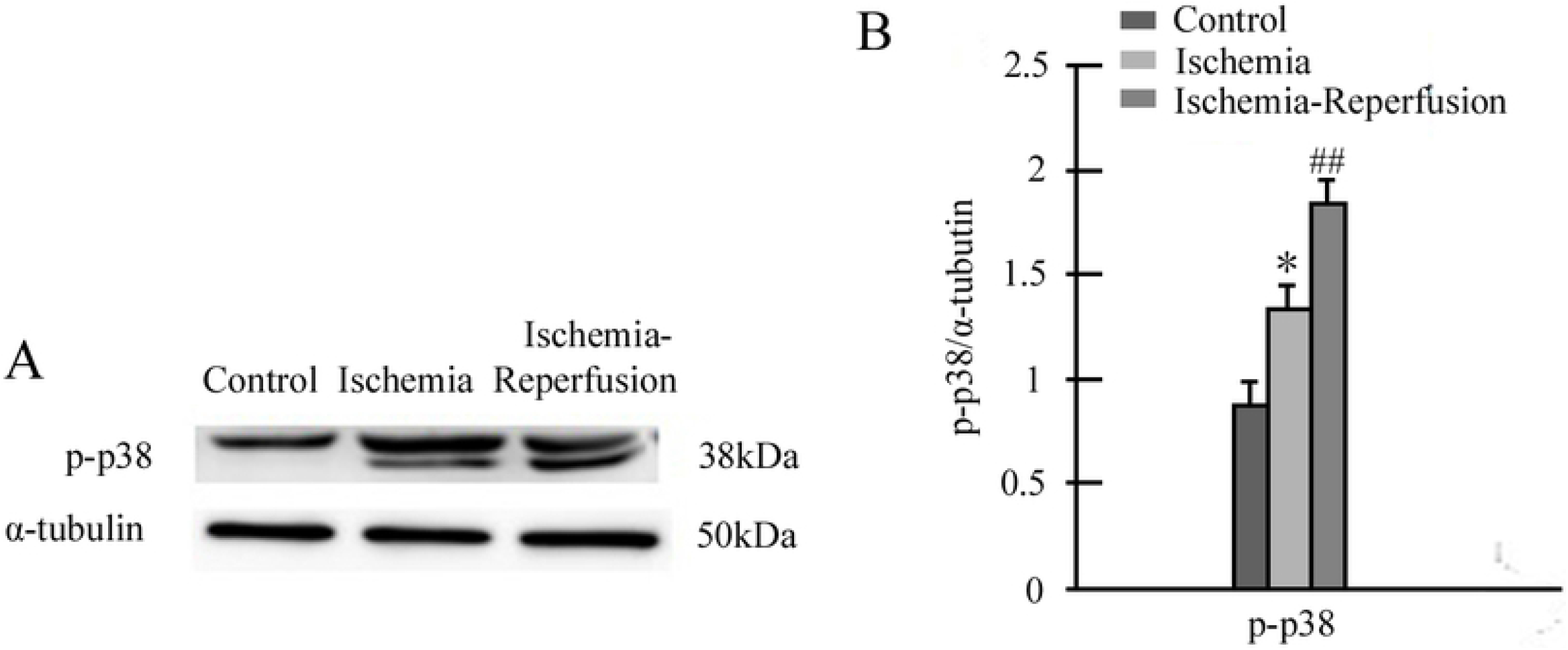
Changes of p-p38 expression in H9c2 cardiomyocytes. (A) western blotting analysis of p-p38 expression, (B) quantitative analysis of protein expression, n = 3. *p< 0.05 vs. control groups; ##p < 0.01 vs. ischemia group. I/R, ischemia/reperfusion.

### Endogenous MT2A inhibits p38 activity expression in H9c2 cardiomyocytes

In order to detect the effect of MT2A on the active expression of p38, we transfected MT2A into H9c2 cardiomyocytes to construct MT2A overexpression cell lines. The expression of p-p38 was detected. The results showed that there was no statistical difference in the expression of p-p38 between the control and the LV-GFP groups. The expression of p-p38 was significantly decreased in the LV-MT2A groups under I/R condition (p< 0.05; Fig. 5).

**Fig. 5.**
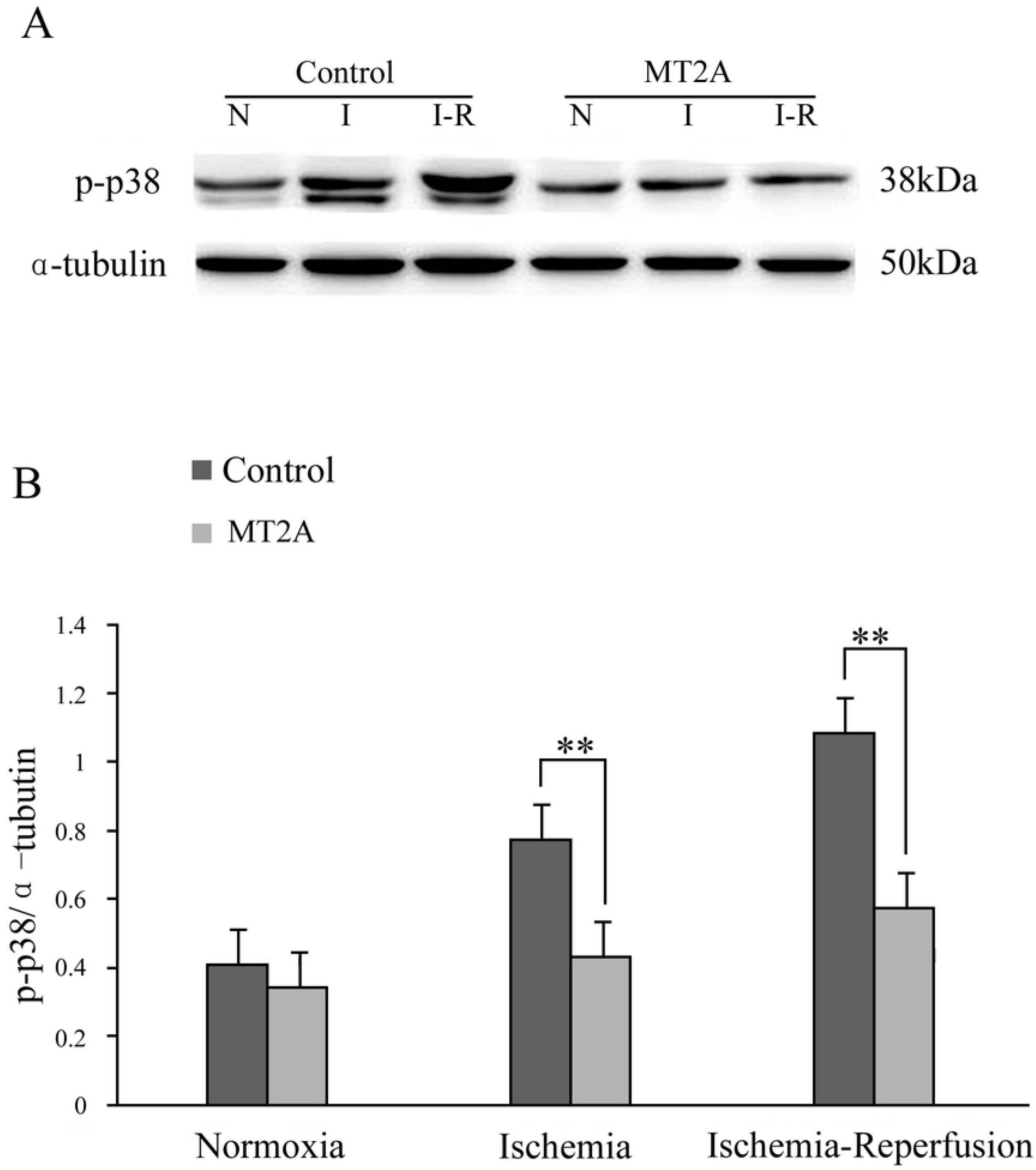
The effect of transfected H9c2 cardiomyocytes on p-p38 was shown. The cells were divided into control, LV-GFP (control vector) and LV-MT2A groups. (A) western blotting analysis of p-p38 protein expression. (B) quantitative analysis of protein expression, n = 3. *p < 0.05 vs. LV-GFP. C: control; V: LV-GFP (control vector); M: LV-MT2A; N, normoxia; I, ischemia; I/R, ischemia/reperfusion.

### Exogenous MT2A inhibits the expression of p-p38 in H9c2 cardiomyocytes

As shown in Fig. 6, MT2A was added to H9c2 cardiomyocytes to observe the expression of p-p38 activity. After adding 10umol/L MT2A, p-p38 expression was significantly inhibited (p < 0.01). The results showed that exogenous addition of MT2A inhibits the expression of p38 activity in H9c2 cardiomyocytes.

**Fig. 6.** The effect of exogenous MT2A on p38 in H9c2 cardiomyocytes was shown. The H9c2 cardiomyocytes were divided into control (Control) and MT2A groups (MT2A). (A) western blotting analysis of p-p38 protein expression. (B) quantitative analysis of protein expression, n = 3. *p < 0.05 vs. control groups. N, normoxia; I, ischemia; I/R, ischemia/reperfusion.

## Discussion

According to the existing research results, the mechanism of myocardial I/R injury is complicated, which is affected by a variety of factors, including metabolic disorders, oxygen free radical, inflammatory response and calcium overload [17]. The pathological changes caused by myocardial I/R injury mainly include the fracture, disordered arrangement and enlarged gap of myocardial fibers, and myocardial dissolution and necrosis, which can lead to heart failure, cardiac dysfunction and abnormal function of myocardial cells [19]. The cardiac function of patients will be greatly reduced and the clinical therapeutic effects (such as bypass surgery, interventional therapy and thrombolysis) will be limited once I/R occurs [20]. Our study has demonstrated that the number of apoptotic cells increased significantly under I/R. It can be seen from the results that the apoptotic pathway mediated by caspase-3 can be activated under ischemia and further activated by I/R. The expression of p-p38 was significantly increased after I/R. This suggested that p-p38 was activated under I/R. We found that both endogenous overexpressed MT2A and exogenously added MT2A can inhibit the active expression of p38 during I/R, suggesting that MT2A intervention can effectively inhibit p38 phosphorylation and decrease the activity of p38 and that p38 is one of the molecules of MT2A against I/R injury in cardiomyocytes.

Apoptosis, a form of programmed cell death, occurs in a wide range of physiological and pathological situations [21]. It is characterized by cell shrinkage, programmed DNA degradation, and increased expression of caspase-3 [22]. Caspase-3 has been implicated as the key protease executing the apoptotic cascade in the intrinsic (mitochondrial) and extrinsic (death ligand) apoptotic pathways [23-25]. In this study, we established a model of I/R in vitro. Our results illustrated that caspase-3 increased significantly under I/R. I/R can further aggravate the apoptosis of H9c2 cardiomyocytes induced by ischemia, which were supported by the results of our previous studies [26] and Xie et al [27].

The expression of p-p38 was significantly increased under I/R (p<0.01). Our results showed that the active expression of p38 was promoted by I/R. The p38 MAPK playing an indispensable role in cell signal transduction, which can cause inflammation, cell growth, apoptosis and other biological effects [4, 5]. The p38 MAPK pathway can be activated during cell ischemia and hypoxia [28]. Yan et al. [29] and Xie et al. [27] reported that the activation of p38 MAPK pathway can induce cardiomyocyte apoptosis and caspase-3 protein expression in H9c2 cells during I/R, thereby aggravating the myocardial I/R injury.

The results of this study indicated that the expression of p-p38 in I/R was significantly increased and decreased significantly after MT2A treatment, we found that both endogenously overexpressed MT2A and exogenously added MT2A can inhibit the active expression of p38 during I/R, suggesting that MT2A intervention can effectively inhibit p38 phosphorylation and decrease the activity of p38 and that p38 is one of the molecules of MT2A against I/R injury in cardiomyocytes. The function of MT2A is to regulate metal homeostasis, detoxification, oxidative stress, immune defense, cell cycle progression, cell proliferation and differentiation, and angiogenesis [30, 31]. Over expression of MT2A can decrease oxygen consumption, down-regulate cellular ATP levels and decrease oxidative phosphorylation capacity, and interact with mitochondrial complexes indirectly, which might be involved in the inhibition of certain respiratory enzymes via metal binding [32]. MT2A could suppress I/R-induced myocardial apoptosis mediated by mitochondrial stress [33]. MT2A is a protective protein from apoptosis by down-regulating the expression of Bax, caspase-3, caspase-9, and caspase-12 [13]. A great deal of evidence shows that oxidative stress is the key mediator of myocardial injury during I/R [34]. Therefore, it is necessary to study and explore the endogenous mechanism or exogenous intervention to inhibit the myocardial injury caused by oxidative stress and apoptosis in the process of I/R. Inhibiting the p38 MAPK pathway attenuated the I/R-induced increases in H9c2 cell apoptosis and caspase-3 protein expression [29].

As far as we know, this study demonstrated that MT2A can significantly reduce the apoptosis of H9c2 cardiomyocytes induced by I/R through p38 MAPK pathway. This study is helpful for us to better understand the protective mechanism of MT2A on I/R.

### Study limitation and future directions

This limitation of the present study means that further research is required. Myocardial I/R injury should be paid attention to achieve a successful clinical reperfusion therapy, such as percutaneous coronary intervention, thrombolysis, coronary artery bypass surgery. A number of mechanisms have been proposed to explain the mediation of reperfusion injury. These include cellular calcium loading, endothelium cell swelling, impaired vascular relaxation, and the action of oxygen radicals [35]. In clinical situations, the outcomes for acute coronary syndrome patients who are suffering from I/R are influenced by many unpredictable events, such as myocardial stunning, coronary vascular spasms, or lethal arrhythmia [36]. The molecular signaling changes of myocardial I/R injury which are needed to be further studied. In addition, the results derived from H9c2 cells may not represent the same mechanisms as in primary cardiomyocytes. More in-depth understanding of the functional, biochemical and molecular characteristics of MT2A will bring a promising novel therapeutic approach for I/R injury. Novel treatments that inhibit or ameliorate the disease process must be developed to allow sufficient time for subsequent treatment.

## Conclusions

We found that MT2A inhibits p38 activity expression, protects cardiomyocytes from I/R injury.

## Acknowledgments

The authors would like to acknowledge Qingmin Feng for her assistance with aspects of the in vitro culturing of the cells. And, we also thank the anonymous reviewers for their helpful suggestions in editing this manuscript.

## Author Contributions

Conceptualization: Jike Li.

Data curation: Ying Zhao, Xiaoli Huang, Yuanlin Lei, and Jike Li.

Formal analysis: Ying Zhao, Xiaoli Huang, Yuanlin Lei, and Jike Li.

Funding acquisition: Jike Li.

Investigation: Jike Li.

Methodology: Ying Zhao, Xiaoli Huang, Yuanlin Lei, and Jike Li.

Project administration: Jike Li.

Supervision: Jike Li.

Validation: Jike Li.

Writing – original draft: Ying Zhao, Xiaoli Huang, Yuanlin Lei, and Jike Li.

Writing – review & editing: Ying Zhao, Xiaoli Huang, Yuanlin Lei, and Jike Li.

